# Role of Kir4.1/Kir5.1 in mediating Angiotensin-II (Ang-II)-induced stimulation of thiazide-sensitive Na-Cl cotransporter

**DOI:** 10.1101/2024.07.19.604336

**Authors:** Xin-Peng Duan, Xin-Xin Meng, Yu Xiao, Cheng-Biao Zhang, Ruimin Gu, Dao-Hong Lin, Wen-Hui Wang

**Author notes:** Corresponding address Dr. Wen-Hui Wang, Department of Pharmacology New York Medical College Valhalla, NY 10595. Two authors contribute equally to the work (Xin-Peng Duan, Xin-Xin Meng).

## Abstract

**Background:** Angiotensin-II (Ang-II) perfusion stimulates Kir4.1/Kir5.1 in the distal-convoluted-tubule (DCT) and thiazide-sensitive Na-Cl-cotransporter (NCC). However, the role of Kir4.1/Kir5.1 in mediating the effect of Ang-II on NCC is not understood.

**Methods:** We used immunoblotting and patch-clamp-experiments to examine whether Ang-II-induced stimulation of NCC is achieved by activation of Kir4.1/Kir5.1 of the DCT using kidney-renal-tubule-specific AT1aR-knockout (Ks-AT1aR-KO), Ks-Kir4.1-knockout and the corresponding wild-type mice.

**Results:** Ang-II perfusion for 1, 3 and 7 days progressively increased phosphor-NCC (pNCC) and total-NCC (tNCC) expression and the effect of Ang-II-perfusion on pNCC and tNCC was abolished in Ks-AT1aR-KO. Ang-II perfusion for 1-day robustly stimulates Kir4.1/Kir5.1 in the late DCT (DCT2) and to a lesser degree in the early DCT (DCT1), an effect was absent in Ks-AT1aR-KO mice. However, Ang-II perfusion for 7-days did not further stimulate Kir4.1/Kir5.1 in the DCT2 and only modestly increased Kir4.1/Kir5.1-mediated K^+^ currents in DCT1. Deletion of Kir4.1 not only significantly decreased the expression of pNCC and tNCC but also abolished the effect of 1-day Ang-II perfusion on the expression of phospho-with-no-lysine-kinase-4 (pWNK4), phosphor-ste-20-proline-alanine-rich-kinase (pSPAK), pNCC and tNCC. However, 7-days Ang-II perfusion was still able to significantly stimulate the expression of pSPAK, pWNK4, pNCC and tNCC, and increased thiazide-induced natriuresis in kidney-tubule-specific Kir4.1 knockout (Ks-Kir4.1 KO) mice without obvious changes in K^+^ channel activity in the DCT.

**Conclusions:** Short-term Ang-II induced stimulation of pWNK4, pSPAK and pNCC depends on Kir4.1/Kir5.1 activity. However, long-term Ang-II is able to directly stimulate pWNK4, pSPAK and pNCC by a Kir4.1/Kir5.1 independent mechanism.

## Introduction

Thiazide-sensitive Na-Cl-cotransporter (NCC) encoded by *Slc12A3* is expressed in the apical membrane of the distal convoluted tubule (DCT)^1–4^. NCC is responsible not only for reabsorption of 5-9% filtered Na^+^ load but also plays a key role in regulating renal K^+^ excretion in the aldosterone-sensitive-distal nephron (ASDN) by controlling Na^+^ delivery to the ASDN^5–7^. For instance, increased dietary K^+^ intake has been shown to inhibit NCC activity thereby increasing Na^+^ delivery to the ASDN ^8, 9^. The increased Na^+^ delivery is expected to enhance ENaC-dependent K^+^ excretion in the ASDN ^10^. In contrast, Na^+^-restriction has been shown to stimulate NCC activity thereby enhancing Na^+^ absorption in the DCT and decreasing Na^+^ delivery to the ASDN ^8, 11^. Thus, the regulation of NCC activity not only has an effect on blood pressure but also plays a role in maintaining K^+^ homeostasis. Ang-II has been shown to stimulate NCC by modulating with-no-lysine kinase (WNK) activity^12–14^, which is the upstream protein kinase for phosphorylating SPAK thereby stimulating its activity^15–18^. The basolateral Kir4.1/Kir5.1 channel activity of the DCT plays a role in the regulation of WNK activity by affecting the intracellular Cl^-^ concentrations^19, 20^. It is well demonstrated that Kir4.1/Kir5.1 heterotetramer in the DCT, a 40 pS inwardly-rectifying K^+^ channel, is the predominate type of basolateral K^+^ channel and responsible for determining the negativity of DCT membrane potential ^21–24^. We have further demonstrated that the Kir4.1/Kir5.1 activity of the DCT is closely related to the NCC activity/expression such that high Kir4.1/Kir5.1 activity is associated with increased NCC activity/expression whereas low Kir4.1/Kir5.1 activity results in decreased NCC activity/expression ^9, 11, 25, 26^. We have previously demonstrated that Ang-II is able to acutely stimulate the basolateral Kir4.1/Kir5.1 in the DCT2 and that prolonged application of Ang-II also stimulates Kir4.1/Kir5.1 in the DCT1 ^27, 28^. This raises a question whether Ang-II-induced stimulation of Kir4.1/Kir5.1 activity is involved in mediating the effect of Ang-II on NCC activity/expression. Thus, the aim of the present study is to examine the role of Kir4.1/Kir5.1 channel activity in the DCT in mediating the effect of Ang-II on NCC activity/expression.

## Methods

### Material and Methods

All supporting data and detailed methods including animal preparation, electrophysiology and immunoblotting are available within the article and its online supplementary file.

### Animals

We used male (m) and female (f) C57/BL6j mice, *Agtr1a^flox/flox^* mice, *Kcnj10^flox/flox^* mice, kidney-tubule-specific AT1aR KO (Ks-AT1aR KO) mice and Ks-Kir4.1 KO mice for the experiments. C57/BL6j mice were purchased from Jackson Lab or the second affiliated hospital animal facility of Harbin Medical University. All transgenic mice have C57/BL6j background and have been bred in New York Medical College animal facility. To generate Ks-AT1aR KO mice, mice expressing Pax8-rtTA and tet-on LC-1 transgene were crossed with *Agtr1a*-floxed mice (a gift from Susan Gurley’s lab of Oregon Health & Science University) to generate inducible Ks-AT1aR knockout mice ^29^. To generate Ks-Kir4.1 KO mice, mice expressing Pax8-rtTA and tet-on LC-1 transgene were crossed with *Kcnj10*-floxed mice. *Agtr1a* or *Kcnj10* gene deletion was conducted in 8-week-old m/f mice homozygous for floxed *Agtr1a* or floxed *Kcnj10* gene and heterozygous for Pax8-rtTA/LC-1 transgene by providing doxycycline (5mg/ml, 5% sucrose) in the drinking water for 2 weeks. This was followed by at least 2 additional weeks without doxycycline treatment before performing experiments. Littermate mice of the same age and genetic background drinking 5% sucrose were used as controls. Tail DNA was PCR amplified and the primers for genotyping are shown in table s1. We harvested the kidney for immunoblotting and electrophysiological studies. The method for obtaining the kidney is described in the supplementary material. The animal use protocol has been reviewed and approved by either NYMC IACUC or Harbin Medical University Animal use committee.

### Patch-clamp experiments

A Narishige electrode puller (Narishige, Japan) was used to make the patch-clamp pipettes from Borosilicate glass (1.7-mm OD). The resistance of the pipette was 2 MΩ when it was filled with solution containing (in mM) 140 KCl, 1.8 MgCl_2_ and 10 HEPES (titrated with KOH to pH 7.4). *Measurement of Kir4.1/Kir5.1-mediated K^+^ currents* We have used the whole-cell recording to measure Ba^2+^-sensitive Kir4.1/Kir5.1-mediated K^+^ currents in the DCT. An Axon 200A patch-clamp amplifier was used to record the whole-cell K^+^ currents, which were low-pass filtered at 1 KHz, digitized by an Axon interface (Digidata 1440A). The pipette solution contains (in mM) 140 KCl, 2 MgCl_2_, 1 EGTA and 5 HEPES (pH 7.4) with 0.5 mM MgATP, whereas the bath solution contains (in mM) 140 KCl, 1.8 CaCl_2_, 1.8 MgCl_2_, and 10 HEPES (pH 7.4). For the measurement of the Kir4.1/Kir5.1 mediated K^+^ currents in the DCT2, 400 nM tertiapin Q (TPNQ, Sigma) was added into split-open DCT to block the ROMK-mediated K^+^ currents ^30^. The whole-cell K^+^ current was determined by adding 1 mM Ba^2+^ in the bath solution. Data were analyzed using the pClamp Software System 9 (Axon).

#### Measurement of inward-to-outward current reversal potential (I-reversal potential)

For measuring I-reversal potential of the DCT, an index of cell membrane potential, the isolated DCT was super-fused with a bath solution containing (in mM) 140 NaCl, 5 KCl, 1.8 CaCl_2_, 1.8 MgCl_2_, and 10 HEPES (pH 7.4) . The pipette was filled with 140 mM KCl pipette solution and was then backfilled with amphotericin B (20 mg/0.1ml). After forming a high-resistance seal (>2 GΩ), the membrane capacitance was monitored until the whole-cell patch configuration was formed. The voltage at which the inward currents were zero was the I - reversal potential determined by a ramp protocol from -100 to 100 mV.

#### Single-channel recording

An Axon 200B patch-clamp amplifier was used to record the single K^+^ channel currents, which were low-pass filtered at 1 KHz and digitized by an Axon interface (Digidata 1440A). Channel activity, defined as NP_o_ (a product of channel number and open probability), was calculated from data samples of 60 seconds duration in the steady state as follows:

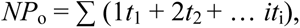

where *t_i_* is the fractional open time spent at each of the observed current levels. The bath solution contains (in mM) 140 NaCl, 5 KCl, 1.8 CaCl_2_, 1.8 MgCl_2_, and 10 HEPES (pH 7.4) and the pipette solution contains (in mM) 140 KCl, 2 MgCl_2_, 1 EGTA and 5 HEPES (pH 7.4). Data were analyzed using the pCLAMP Software System 9 (Axon).

### Immunoblotting

Renal cortex protein extract was obtained from frozen tissue samples homogenized in a buffer containing 250 mM sucrose, 50 mM Tris-HCl (pH 7.5), 1 mM EDTA, 1mM EGTA, 1 mM DTT supplemented with phosphatase and protease inhibitor cocktails (Sigma). Protein (40-60 μg) was separated on 4-12% (wt/vol) Tris-Glycine gel (Novex^TM^, TheromoFisher Scientific) and transferred to nitrocellulose membrane. The membranes were incubated for 1 hour with LI-COR blocking buffer (PBS) and then incubated overnight at 4°C with primary antibodies including anti-NCC, anti-pNCC at threonine-53, anti-Kir4.1, anti-pWNK4, anti-WNK4, anti-pSPAK and anti-SPAK . All antibodies used in the experiments are validated and the information is included in table s2. An Odyssey infrared imaging system (LI-COR) was used to capture the images at a wave-length of 680 or 800 nM.

### Statistical analysis

We used software (Sigma plot 14) for the statistical analysis. For analyzing the values between two groups we used t-test, and for comparisons of the values within the same group, we used paired t-test. We used one-way or two-way ANOVA for analyzing results of more than two groups, and Holm-Sidak test was used as post-hoc analysis. *P*-values <0.05 were considered statistically significant. Data are presented as the mean ± SEM.

## Results

To examine the effect of Ang-II on the expression of pNCC and tNCC, m/f control mice were continuously perfused at a no-pressure dose of Ang-II (200 ng/min/Kg body-weight) or vehicle through an osmotic pump for 1-day, 3-days and 7-days, respectively. The renal tissues were then harvested for immunoblotting or for electrophysiology experiments. Fig. 1A is a set of western blots showing the expression of pNCC and tNCC in m/f control mice (Fg.s1 shows uncut westerb blot) and Fig. 1B is a bar graph with scatter plots summarizing each data point and the mean value of the normalized band density. In comparison to vehicle-treated animals (0-day Ang-II), Ang-II perfusion for 1-, 3- and 7-days progressively increased the expression of pNCC in male mice by 52±4%, 96±8% and 163±14% (n=6), and in female mice by 48±4%, 100±9% and 155±14% (n=6), respectively. Ang-II perfusion for 1-, 3- and 7-days also progressively increased the expression of tNCC in male mice by 40±3%, 85±8 % and 135±8%, and in female mice by 48±4%, 96±8% and 160±13% in comparison to vehicle. To examine whether the effect of Ang-II on NCC depends on AT1aR, we repeated the experiments in Ks-AT1aR KO mice and control mice (*Agtr1a^flox/flox^*). Fig 1C is a set of western blot showing the expression of pNCC and tNCC in male Ks-AT1aR KO and *Agtr1a^flox/flox^*mice (Fig.s2A shows uncut western blot) and Fig. 1D is a bar graph with scatter plots summarizing each data point and the mean value of the normalized band density. It is apparent that deletion of AT1aR completely abolished the stimulatory effect of Ang-II on pNCC and tNCC, indicating that the effect of Ang-II on NCC was mediated by AT1aR.

**Fig 1.**
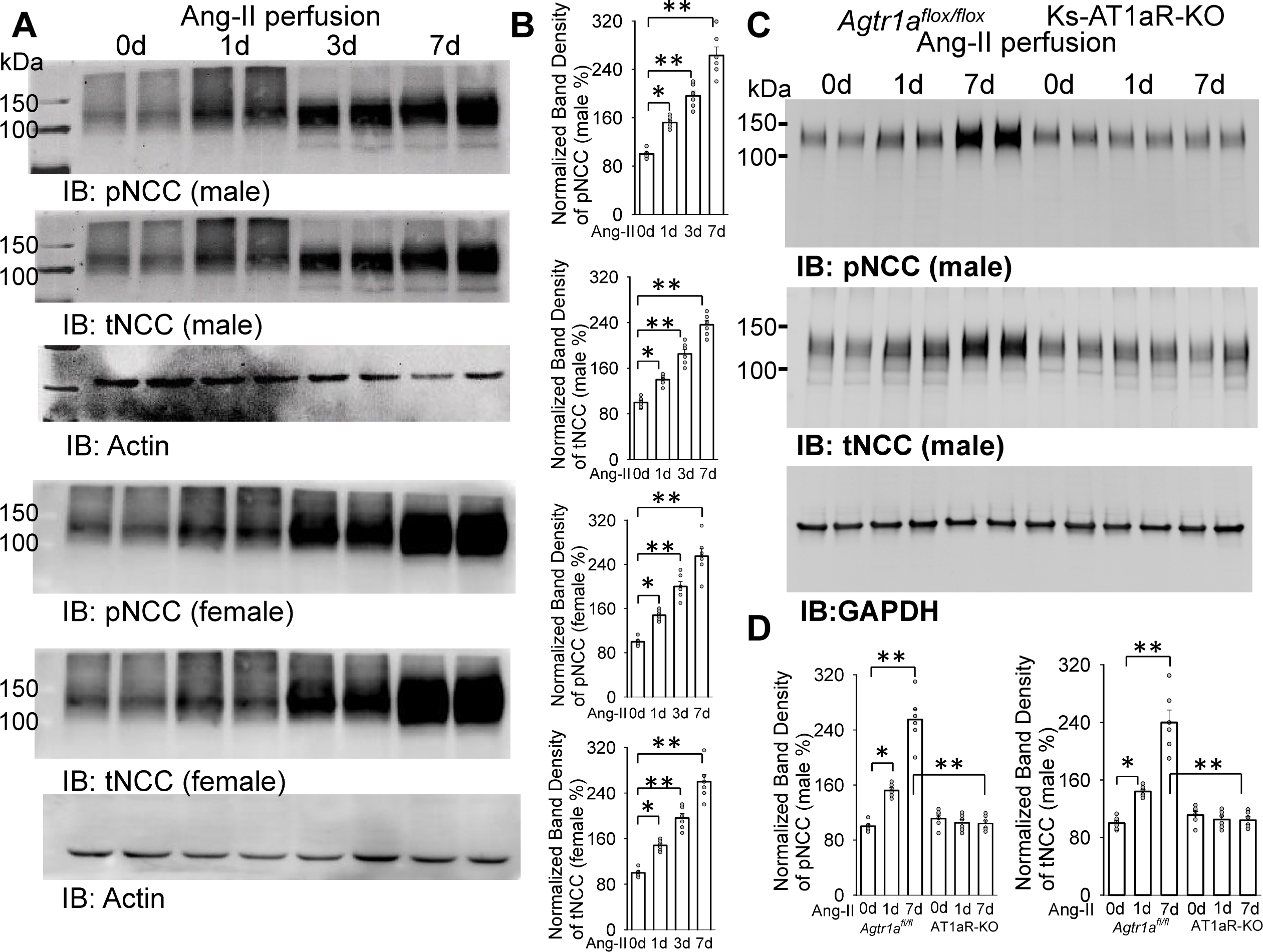
Ang-II perfusion stimulates the expression of NCC via AT1aR. (A) A set of western blot shows the expression of phosphor (p)-NCC (Thr^53^), total NCC and actin in male and female control mice (C57/BL6j) treated with vehicle (0-day Ang-II) or Ang-II at no pressure dose (200 ng/min/Kg BW, via osmotic pump) for 1 day (1d), 3d and 7d. (B) A bar graph with scatter plots summarizes mean value (n=6) and each data point of the normalized band density of pNCC or tNCC for m/f mice. (C) A western blot shows the expression of pNCC and tNCC in male *Agtr1a^flox/flox^*mice or Ks-AT1aR KO mice treated with vehicle (0-day Ang-II) or Ang-II for 1d and 7d. (D) A bar graph with scatter plots summarizes mean value (n=6) and each data point of the normalized band density of pNCC or tNCC. Single asterisk and double-asterisks indicate p-value<0.05 and p-value<0.01, respectively.

Our previous study demonstrated that Ang-II perfusion for 7 days stimulated the basolateral Kir4.1/Kir5.1 in the DCT, an effect was absent in Ks-AT1aR KO mice^27^. To determine whether Ang-II also progressively stimulates Kir4.1/Kir5.1, we next examined the Kir4.1/Kir5.1-mediated K^+^ currents in the DCT2 of male control mice treated with vehicle or Ang-II for 1-day and 7-days. The reason for examining the effect of Ang-II on Kir4.1/Kir5.1 in DCT2 is that Ang-II has been shown to initially target Kir4.1/Kir5.1 of DCT2 via AT1aR ^28^. Fig. 2A is a set of traces demonstrating Kir4.1/Kir5.1-mediated K^+^ currents in the DCT2 of male *Agtr1a*^flox/flox^ and Ks-AT1aR KO mice measured with whole-cell voltage-clamp using step-protocol from -100 to 60 mV. Fig.2B is a scatter plot summarizing each data point and mean value of whole-cell Kir4.1/Kir5.1-mediated K^+^ currents at -60 mV. Ang-II perfusion for 1-day robustly increased Kir4.1/Kir5.1-mediated K^+^ currents from 56±2 pA/pF (vehicle) to 104±6 pA/pF (n=6) in *Agtr1a*^flox/flox^ mice. However, Ang-II perfusion for 7-days did not further increase Kir4.1/Kir5.1-mediated K^+^ currents in DCT2 (101±6 pA/pF). Thus, prolonged Ang-II perfusion did not progressively stimulate Kir4.1/Kir5.1. This notion was also supported by the experiments in which the membrane potential of DCT2 was measured. Fig. 2C is a set of traces showing I-reversal-potential of the DCT2 in male *Agtr1a*^flox/flox^ and Ks-AT1aR KO mice measured with the whole-cell-voltage-clamp. Fig.2D is a scatter plot summarizing each data point and mean value of I-reversal-potential of the DCT2 in the mice treated with vehicle or Ang-II for 1 and 7-days. It is apparent that Ang-II perfusion for 1-day increased the negativity of I-reversal-potential in DCT2 (from -61±1 to -72±1 mV, n=6). However. 7-days Ang-II perfusion did not further hyperpolarize DCT2 membrane because I-reversal-potentials of DCT2 were similar between 1- day and 7-day Ang-II (-73±1 mV). Again, the effect of Ang-II on the DCT membrane potential was absent in AT1aR KO mice. Thus, it is unlikely that Ang-II (7-days) perfusion-induced increase in NCC expression was due to progressively stimulating Kir4.1/Kir5.1 in the DCT2.

**Fig 2.**
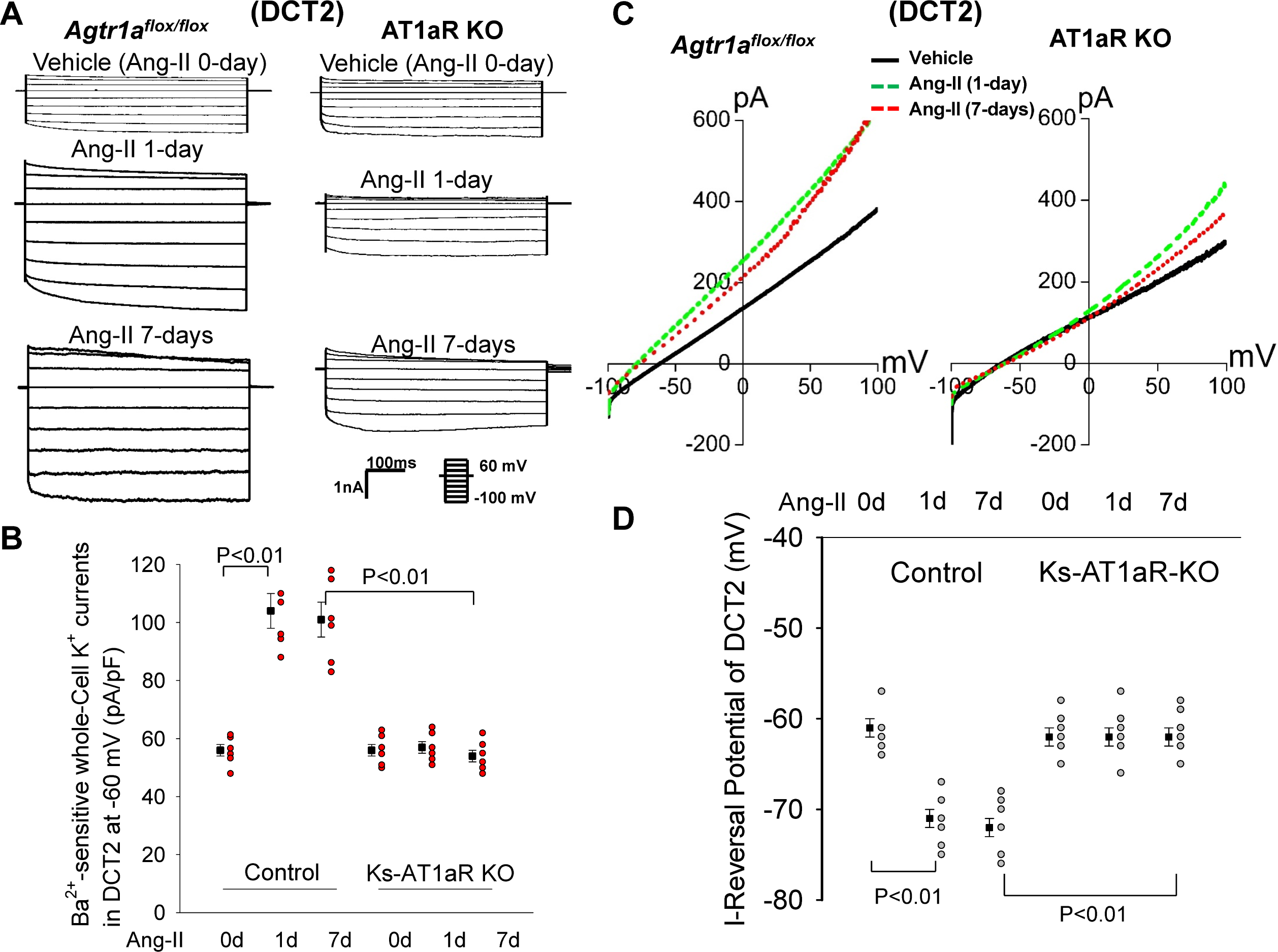
Ang-II perfusion stimulates Kir4.1/Kir5.1 of DCT2 via AT1aR. (A) A set of traces shows whole-cell Ba^2+^-sensitive Kir4.1/Kir5.1 mediated-K^+^ currents using step-protocol from - 100 to 60 mV in DCT2 of male *Agtr1a^flox/flox^* mice and Ks-AT1aR KO mice treated with vehicle or Ang-II for 1d and 7d. (B) The above experiments are summarized in a scatter plot showing the mean value (n=6) and each data point measured with whole-cell recording at -60 mV in DCT2. (C) A set of traces of I-reversal potential measured with whole-cell voltage clamp from -100 to 100 mV in the DCT2 of male *Agtr1a^flox/flox^* mice and Ks-AT1aR KO mice treated with vehicle (black), 1-day-Ang-II (green) and 7-days-Ang-II (red). (D) A scatter plot summarizes the experiments in which I-reversal potential was measured in DCT2 of male AT1aR KO mice and *Agtr1a^flox/flox^* mice. The mean value (n=6) of each group is shown on the left of each column. The significance is determined by two-way ANOVA.

We next examine the possibility that progressive stimulatory effect of Ang-II on NCC may be due to the stimulation of Kir4.1/Kir5.1 of the DCT1. This speculation is based on our previous observation that Ang-II perfusion for 7 days also stimulated the Kir4.1/Kir5.1 in DCT1 via AT1aR ^27^, although Ang-II failed to acutely stimulate Kir4.1/Kir5.1 in DCT1^28^. Thus, we have first used the single channel recording to examine the 40-pS K^+^ channel activity (Kir4.1/Kir5.1) in DCT1 of m/f control mice treated with vehicle or Ang-II for 1-, 3- and 7-days. From the inspection of Fig.3A, it is apparent that Ang-II perfusion for 1-day stimulated the 40- pS K^+^ channel of the DCT1. Fig. 3B is a scatter plot summarizing each data point and mean values of the experiments in which we calculated 40-pS K^+^ channel activity, defined by NP_o_, in the DCT1 of m/f mice treated with Ang-II for 1, 3, 7-days, respectively. Ang-II for 1-day increased NP_o_ of the 40-pS K^+^ channel from 1.19±0.04 (vehicle) to 2.10±0.07 in 6 male mice and from 1.20±0.05 (vehicle) to 2.2±0.06 in 6 female mice. However, Ang-II perfusion for 3 and 7-days tends to slightly increase 40-pS K^+^ channel NP_o_ in m/f mice but the effect was not significant in comparison to 1-day Ang-II. We have also used the whole-cell recording to measure Kir4.1/Kir5.1-mediated whole-cell K^+^ currents in DCT1 of male control mice treated with vehicle or Ang-II for 1 and 7-days. Ang-II perfusion for 1-day increased Kir4.1/Kir5.1- mediated K^+^ current from 54±3 pA/pF (vehicle) to 86±2 pA/pF (n=6). However, prolonged Ang- II perfusion only modestly increased the K^+^ currents to 98±4 pA/pF (n=6). Thus, it is also unlikely that long-term effect of Ang-II perfusion on NCC expression was due to stimulating Kir4.1/Kir5.1 in the DCT1.

**Fig 3.**
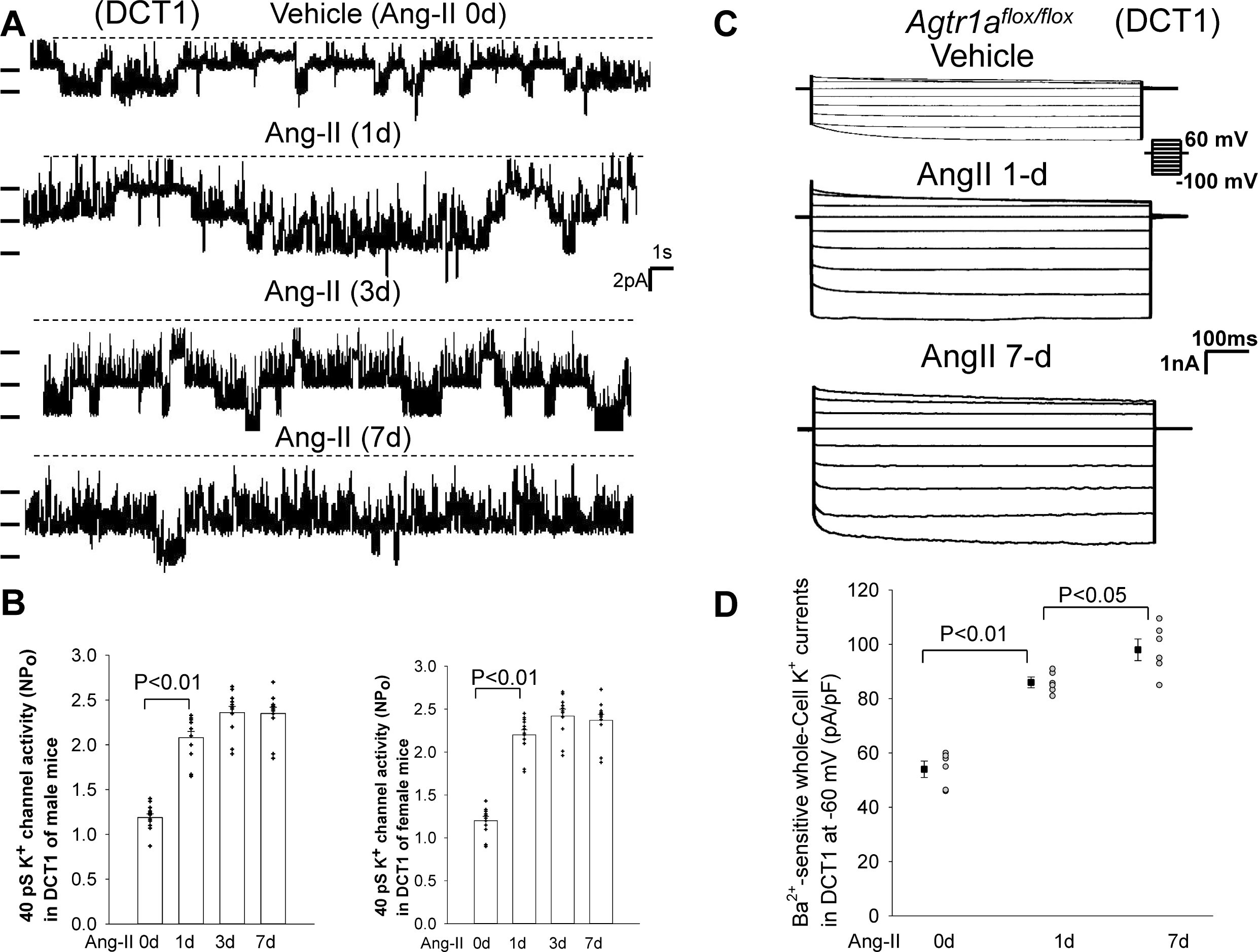
Kir4.1/Kir5.1 activity of DCT1 has reached a submaximal level after 1d Ang-II. (A) A single channel recording made in a cell-attached patch shows the 40 pS K^+^ channel activity in the DCT1 of male control mice treated with vehicle (0 day Ang-II) or Ang-II for 1 day (1d), 3d and 7d. (B) The bar graph with scatter plot showing the effect of Ang-II on the 40 pS K^+^ channel activity, defined by NP_o_, in m/f mice treated with vehicle (0 day Ang-II), Ang-II for 1d, 3d and 7d. (C) A set of traces shows whole-cell Ba^2+^-sensitive Kir4.1/Kir5.1 mediated-K^+^ currents using step-protocol from -100 to 60 mV in DCT1 of male *Agtr1a^flox/flox^* mice treated with vehicle or Ang-II for 1 day and 7 days. (D) The above experiments are summarized in a scatter plot showing the mean value (n=6) and each data point measured with whole-cell recording at -60 mV in DCT2. The mean value of each group is shown in the left of each column.

Because 7-day Ang-II perfusion-induced stimulation of NCC was not associated with further increase in Kir4.1/Kir5.1 activity, we speculate whether Ang-II may also stimulate NCC activity/expression by Kir4.1/Kir5.1-indepent mechanism. Thus, we next examined the effect of Ang-II on NCC in male Ks-Kir4.1-KO mice. Fig.4A is a western blot showing the expression of pNCC and tNCC in male *Kcnj10^flox/flox^* mice and Ks-Kir4.1 KO mice treated with vehicle and Ang-II perfusion for 1 or 7 days (Fig.s2B shows uncut western blots) and Fig. 4B is a bar graph with scatter plot showing the normalized band density of the experiments. From inspection of Fig. 4, it is apparent that Ang-II perfusion for 1-day increased the pNCC expression by 55±6% (n=5) and tNCC expression by 25±6% in comparison to vehicle treated mice. As expected, deletion of Kir4.1 decreased the expression of pNCC and tNCC and also abolished the effect of 1-day Ang-II on NCC. However, 7-days Ang-II perfusion was still able to increase the expression of pNCC from 28±5% of the control value to 52±5% of the control value in Ks- Kir4.1 KO mice, suggesting that Ang-II was able to stimulate NCC by a Kir4.1/Kir5.1 independent mechanism. The notion that 1-day AngII-infusion-induced stimulation of pNCC requires the presence of Kir4.1/Kir5.1 activity in the DCT was also supported by measurement of HCTZ-induced E_Na_. Fig.4C is a scatter plot showing that HCTZ-induced net natriuresis (before and after HCTZ) was larger in the mice treated with 1-day Ang-II (2.71±0.1 μEq/min/100g body-weight, n=5) than vehicle-treated mice (1.21±0.1 μEq/min/100g body-weight). While NCC activity was inhibited in Ks-Kir4.1 KO mice (net HCTZ-induced E_Na,_ 0.18±0.03 μEq/min/100g body-weight), overnight Ang-II treatment had also no significant effect on HCTZ- induced net natriuresis (0.21±0.03 μEq/min/100g body-weight) in Ks-Kir4.1 KO mice, suggesting that Kir4.1 is required for 1-day Ang-II-induced stimulation of NCC. The lack of the overnight Ang-II-induced stimulation of NCC was not due to the strong negative effect of Kir4.1 deletion on NCC since a prolonged AngII perfusion (7 days) was still able to significantly increase HCTZ-induced net natriuresis (0.75±0.06 μEq/min/100g body-weight) in comparison to untreated or overnight Ang-II-treated Ks-Kir4.1 KO mice. Ang-II-induced stimulation of NCC was not related to the DCT membrane potential because 7-days Ang-II perfusion had no effect on DCT2 membrane potential (-40±1 mV, n=4), which was similar to vehicle-treated group in Ks-Kir4.1 KO mice and significantly lower than the control value (-61.4±1 mV, n=7) (Fig.4D). Thus, Ang-II increases NCC expression/activity by Kir4.1-dependent (1-day Ang-II) and Kir4.1- independent mechanisms (7-days Ang-II).

**Fig 4.**
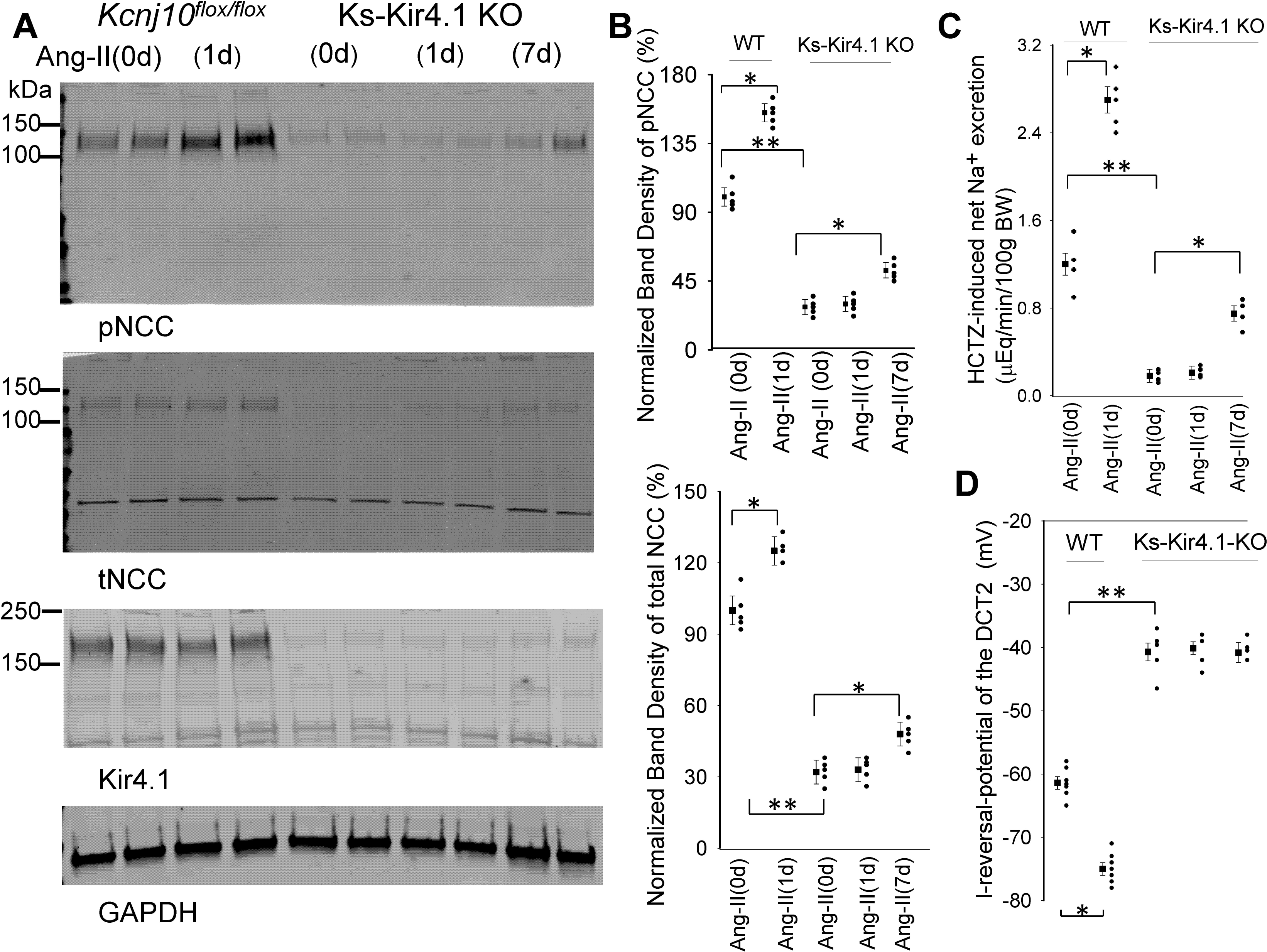
Short-term Ang-II effect on NCC is absent in Ks-Kir4.1 KO mice. (A) A western blot shows the expression of pNCC and tNCC in male *Kcnj10^flox/flox^*mice (n=5) and Ks-Kir4.1 KO mice (n=5) treated with vehicle or Ang-II for 1d and 7d (only in Ks-Kir4.1 KO mice). (B) The above experiments are summarized in a bar graph with scatter plot showing the normalized band density of pNCC and tNCC . (C) A scatter plot shows the mean value (n=4-5) and each data point of the experiments in which HCTZ (30 mg/Kg)-induced natriuresis was measured in male *Kcnj10^flox/flox^* mice and Ks-Kir4.1 KO mice treated with vehicle or Ang-II for 1d and 7d (only in Ks-Kir4.1 KO mice). (D) A scatter plot summarizes the experiments in which I-reversal potential was measured in DCT2 of male *Kcnj10^flox/flox^* and Ks-Kir4.1 KO mice treated with vehicle or Ang-II for 1d and 7d. The mean value of each group (n=4-7) is shown in the left of each column. The significance is determined by two-way ANOVA. Single asterisk and double-asterisks indicate p-value<0.05 and p-value<0.01, respectively.

Because WNK4/SPAK pathway plays a key role in stimulating NCC activity, we next examined the effect of Ang-II on WNK4 and SPAK in male *Kcnj10^flox/flox^* mice and Ks-Kir4.1 KO mice treated with vehicle or Ang-II for 1 and 7 days. Fig.5A is a set of western blots showing the expression of pSPAK, SPAK, pWNK4 at Ser^1160^ (an indication of active WNK4) and total WNK4 (Fig.s3 shows uncut western blots). Fig.5B (for SPAK) and Fig. 5C (for WNK4) are two sets of bar graphs with scatter plots summarizing each data point and mean value of the normalized band density of western blots. Ang-II perfusion for 1-day significantly increased the expression of pSPAK by 37±4% and pWNK4 by 31±3% (n=5) in comparison to vehicle-treated group. However, 1-day Ang-II perfusion failed to increase pSPAK or pWNK4 expression in Ks-Kir4.1 KO mice, suggesting the role of Kir4.1 in mediating the effect of 1-day Ang-II on SPAK and WNK4. In contrast, 7-days Ang-II perfusion was still able to significantly increase pSPAK expression by 74±4%, SPAK expression by 50±4%, pWNK4 expression by 83±4% and WNK4 expression by 40±4% in the Ks-Kir4.1 KO mice, compared to the vehicle-treated group. Moreover, western blot confirmed that Kir4.1 protein was almost completely absent in Ks-Kir4.1 KO mice (Fig.4A). Thus, data indicate that the short-term Ang-II perfusion stimulates SPAK and WNK4 by Kir4.1/Kir5.1 dependent mechanism whereas prolong perfusion of Ang-II perfusion is able to stimulate SPAK and WNK4 by a Kir4.1/Kir5.1 independent mechanism.

**Fig 5.**
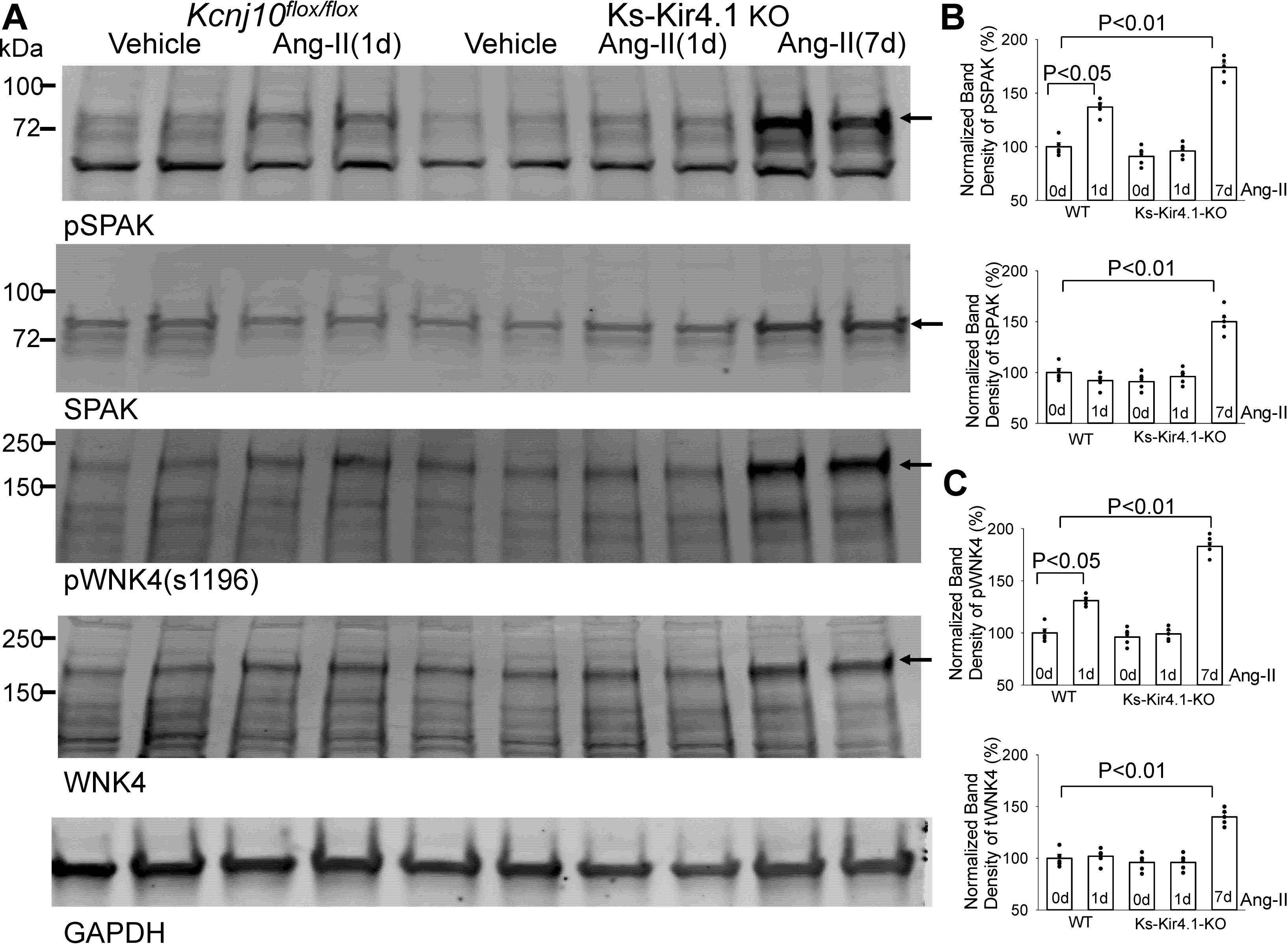
Short-term Ang-II effect on SPAK/WNK4 is absent in Ks-Kir4.1 KO mice. **(A)** A set of western blot shows the expression of pSPAK, SPAK, pWNK4 and WNK4 in male *Kcnj10^flox/flox^* mice (n=5) and Ks-Kir4.1 KO mice (n=5) treated with vehicle or Ang-II for 1d and 7d (only in Ks-Kir4.1 KO mice). (B &C) The above experiments are summarized in a bar graph with scatter plots showing the normalized band density of pSPAK, SPAK, pWNK4 and WNK4. Arrows indicate bands used for the calculation of normalized band density.

## Discussion

The main finding of the present study is to demonstrate the role of Kir4.1/Kir5.1 of the DCT in mediating the effect of short-term Ang-II on thiazide-sensitive NCC. Our present study has confirmed the previous report that Ang-II stimulates NCC expression/activity^13, 31–33^.

However, unlike the previous experiments in which stimulatory effect of Ang-II on NCC expression/activity was observed in the animals treated with Ang-II at one time point ranged from 20 minutes to 8 days ^31, 32^, we have now examined the expression of NCC in the mice received Ang-II perfusion at several time points. We have observed that long-term Ang-II perfusion increased NCC expression more robust than the short-term, although 1-day Ang-II perfusion already significantly increased NCC expression compared to the vehicle-treated animals. Thus, our finding was consistent with previous report that 14-days Ang-II perfusion would increase pNCC expression more robust than 3-days Ang-II perfusion in rats ^34^. Although 7-days Ang-II perfusion increased pNCC more than 1-day Ang-II perfusion, the deletion of AT1aR completely abolished both short-term and long-term effects of Ang-II on pNCC. This strongly suggests the role of AT1aR in mediating the effect of Ang-II on NCC.

Although the effect of Ang-II on NCC is well established, the mechanism by which Ang- II stimulates NCC is not completely understood. It has been reported that 20-minute acute Ang-II treatment stimulates NCC trafficking to the plasma membrane ^14, 31^. Also, Ang-II has been shown to increase phosphorylation of NCC by WNK4-SPAK pathway ^12, 13^. Stimulation of AT1aR by Ang-II activates WNK4 by phosphorylating Ser^1196^ of WNK4, which leads to autophosphorylation of the T-loop of WNK4 thereby activating WNK4 and NCC ^35^. However, it is not explored whether the basolateral Kir4.1/Kir5.1 activity is involved in mediating the effect of Ang-II on WNK4 and NCC. It is well established that the basolateral Kir4.1/Kir5.1 plays an important role in regulating WNK4 activity and NCC expression ^9, 20, 21, 25, 36–38^. We have demonstrated that stimulation of Kir4.1/Kir5.1-induced hyperpolarization is associated with high NCC expression/activity whereas inhibition of Kir4.1/Kir5.1 activity-induced depolarization is correlated with decreased NCC activity/expression ^19^. The hyperpolarization is expected to increase the driving force for the Cl^-^ exit through basolateral membrane thereby decreasing the intracellular Cl^-^ concentrations which should activate the Cl^-^-sensitive WNK activity ^20^. On the other hand, the depolarization of the DCT should decrease the driving force for Cl^-^ exit across the basolateral membrane thereby increasing the intracellular Cl^-^ concentrations and inhibiting WNK4. Indeed, Su et al demonstrated that the inhibition of Kir4.1/Kir5.1 of the DCT raised the intracellular Cl^-^ concentrations ^39^. The possibility that Ang-II may stimulate NCC by activating Kir4.1/Kir5.1 of the DCT is suggested by the finding that Ang-II stimulates the basolateral Kir4.1/Kir5.1 activity.

We have previously demonstrated that acute Ang-II treatment stimulated Kir4.1/Kir5.1 and hyperpolarized DCT2 membrane ^28^, an effect was absent in AT1aR deficient mice, suggesting that the effect of Ang-II on Kir4.1/Kir5.1 was mediated by AT1aR. We have now further demonstrated that 1-day Ang-II perfusion activates Kir4.1/Kir5/1 not only in DCT2 but also in DCT1 to a lesser degree. Three lines of evidence have suggested that short-term Ang-II-induced stimulation of NCC was most likely achieved by activation of Kir4.1/Kir5.1 of the DCT: 1)1-day-Ang-II perfusion stimulated the expression of pNCC and tNCC in the control mice but not in Ks-Kir4.1 KO mice, 2) 1-day-Ang-II treatment increased thiazide-induced natriuresis only in the control mice but not in Ks-Kir4.1 KO mice; 3) 1-day-Ang-II treatment stimulated the expression of pSPAK and pWNK4 only in the control mice but not in the Ks-Kir4.1 KO mice.

Thus, it is conceivable that short-term Ang-II induced stimulation of NCC depends on Kir4.1/Kir5.1 activity. We speculate that acute or short-term Ang-II-induced stimulation of WNK4 and NCC requires the participation of the basolateral Kir4.1/Kir5.1 in the DCT, especially DCT2. Ang-II-induced stimulation of Kir4.1/Kir5.1 hyperpolarizes the DCT membrane thereby activating WNK4 and SPAK which increase NCC expression/activity.

Prolong Ang-II perfusion stimulates NCC expression/activity progressively as evidenced by the fact that 7-days Ang-II-induced stimulation of pNCC was larger than 1-Day Ang-II application. In contrast, prolong Ang-II perfusion did not progressively stimulate Kir4.1/Kir5.1 of the DCT because 7-days Ang-II treatment did not increase Kir4.1/Kir5.1-mediated K^+^ currents in DCT2 in comparison to 1-day Ang-II while it only modestly increases K^+^ currents in DCT1. This finding strongly suggests that prolong Ang-II perfusion stimulates NCC expression/activity by a mechanism other than hyperpolarization of DCT membrane. Three lines of evidence suggest that Ang-II is able to stimulate NCC expression/activity by a Kir4.1/Kir5.1-independent mechanism. First, 7-days-Ang-II perfusion increased the expression of pNCC and tNCC in the Ks-Kir4.1 KO mice. Second, 7-days-Ang-II perfusion enhanced thiazide-induced natriuresis in the Ks-Kir4.1-KO mice. Finally, 7-days-Ang-II perfusion was able to stimulate the expression of pSPAK and pWNK4 in Ks-Kir4.1 KO mice.

### Perspective

It is well established that Ang-II stimulates NCC expression/activity in the DCT. Our present study demonstrates the role of Kir4.1/Kir5.1 in mediating the effect of Ang-II on NCC expression/activity. Fig.6 is a scheme illustrating the mechanism by which Ang-II stimulates NCC by Kir4.1/Kir5.1-dependent mechanism (A) and by K^+^-channel independent mechanism (B). Acute or short-term (less than 24 hours) Ang-II stimulates NCC by a Kir4.1/Kir5.1-dependent mechanism. Presumably, acute or short-term Ang-II activates the Kir4.1/Kir5.1 in the DCT2 but also to a lesser degree in DCT1 thereby hyperpolarizing DCT membrane. The hyperpolarization of DCT membrane is expected to activate Cl^-^ -sensitive WNK-SPAK pathway thereby increasing NCC phosphorylation and activity. However, prolong Ang-II is also able to increase WNK-SPAK pathway which stimulates NCC phosphorylation/activity by Kir4.1/Kir5.1 independent mechanism. One possible mechanism by which Ang-II stimulates NCC by Kir4.1/Kir5.1-independent mechanism is to inhibit degradation mechanism of WNK4 ^40^. It has been shown Ang-II treatment for 4 days increased KLHL3 phosphorylation at serine 433 by PKC. The phosphorylated KLHL3 is not able to bind with WNK4 thereby preventing WNK4 degradation and increasing NCC expression/activity. We conclude that Kir4.1/Kir5.1 activity in the DCT is required for acute and short-term Ang-II induced stimulation of NCC. Prolong Ang- II-treatment is able to stimulate NCC by Kir4.1/Kir5.1-independent mechanism.

**Fig 6.**
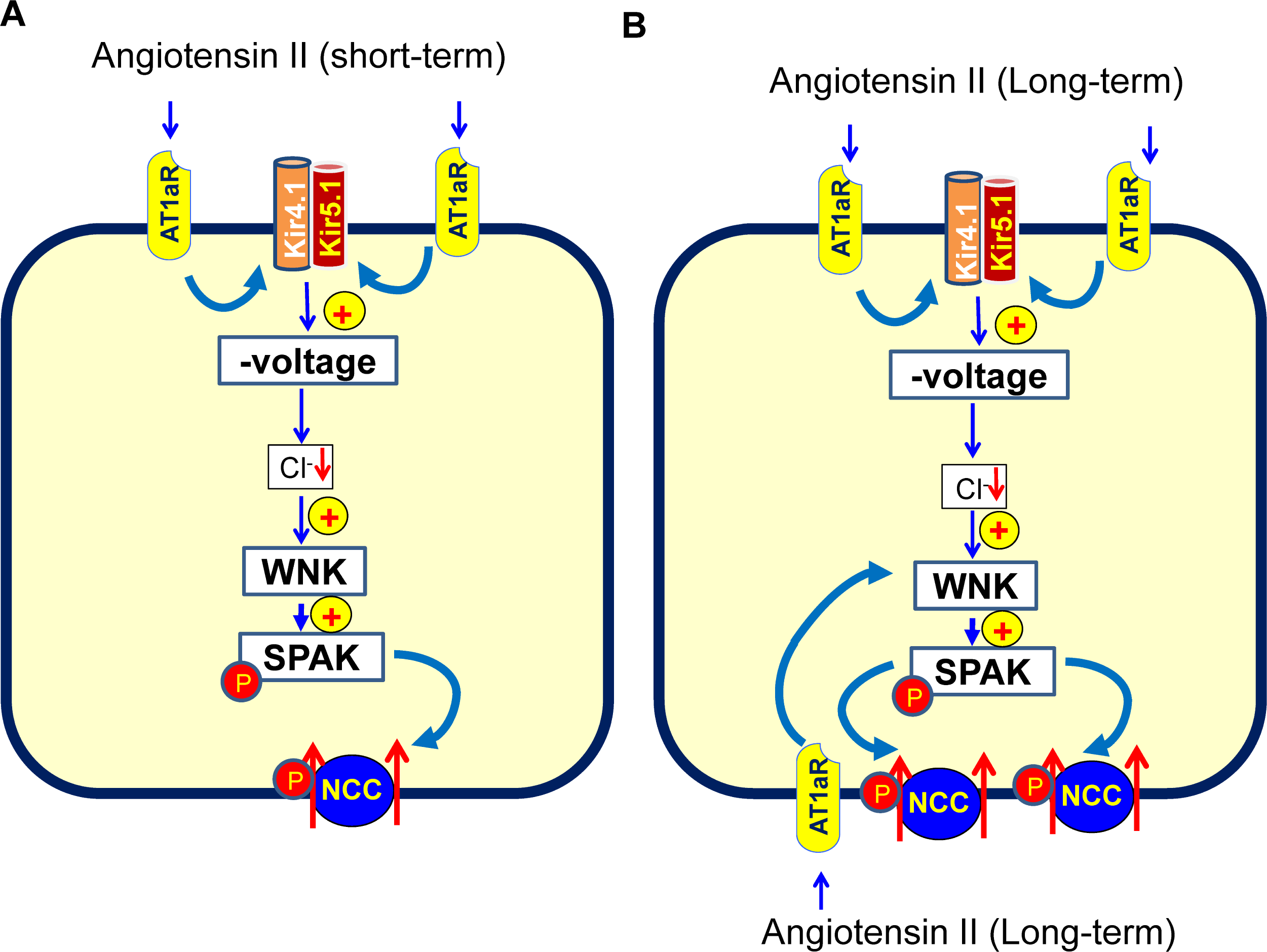
Ang-II stimulates NCC by Kir4.1/Kir5.1-dependent and independent mechanism. A cell scheme illustrating the mechanism by which Ang-II stimulates NCC by Kir4.1/Kir5.1-depemdent (A) and by Kir4.1/Kir5.1-independent mechanism (B). Short-term Ang-II application stimulates the basolateral Kir4.1/Kir5.1 thereby hyperpolarizing DCT membrane which then activates Cl^-^-sensitive WNK. Activated WNK further stimulates NCC phosphorylation by SPAK. Long-term Ang-II may stimulate NCC by Kir4.1/Kir5.1-independent mechanism. The apical location of AT1aR is solely hypothetic.

## Acknowledgments

Authors thank Dr. Susan B Gurley at Oregon Health & Science University) for providing *Agtr1a^flox/flox^* mice.

## NOVELTY and RELEVANCE

### What is New?

- Ang-II perfusion progressively stimulates NCC phosphorylation.
- Deletion of Kir4.1 abolished the effect of short-term Ang-II on WNK, SPAK and NCC.
- A prolong perfusion of Ang-II is required to stimulate WNK4, SPAK and NCC in Kir4.1 deficient mice.

### What is relevant?

- We identify a novel mechanism by which Ang-II stimulates NCC expression/activity by targeting basolateral Kir4.1/Kir5.1 activity of the DCT.
- Acute or short-term Ang-II stimulates NCC, under physiological conditions, by activating Kir4.1/Kir5.1 in the DCT, especially DCT2. Only a sustained Ang-II stimulation is expected to activate WNK-SPAK and NCC by Kir4.1/Kir5.1-independent mechanism.

## Clinical/Pathophysiological implications?

Short-term Ang-II stimulates the basolateral Kir4.1/Kir5.1 of the DCT which then activates Cl^-^- sensitive WNK-SPAK thereby stimulating NCC. Prolonged Ang-II-treatment may stimulate WNK, SPAK and NCC by Kir4.1/Kir5.1-independent mechanism. Thus, inhibiting basolateral K^+^ channel of the distal nephron can effectively block the Ang-II-induced stimulation of kidney sodium absorption.

